# Nonuple atg8 mutant provides genetic evidence for functional specialization of ATG8 isoforms in *Arabidopsis thaliana*

**DOI:** 10.1101/2024.12.10.627464

**Authors:** Alessia Del Chiaro, Nenad Grujic, Jierui Zhao, Ranjith Kumar Papareddy, Peng Gao, Juncai Ma, Christian Lofke, Anuradha Bhattacharya, Ramona Gruetzner, Pierre Bourguet, Frédéric Berger, Byung-Ho Kang, Sylvestre Marillonnet, Yasin Dagdas

**Author notes:** Correspondence: Yasin Dagdas. These authors contributed equally (Alessia Del Chiaro, Nenad Grujic).

## Abstract

Autophagy sustains cellular health by recycling damaged or excess components through autophagosomes. It is mediated by conserved ATG proteins, which coordinate autophagosome biogenesis and selective cargo degradation. Among these, the ubiquitin-like ATG8 protein plays a central role by linking cargo to the growing autophagosomes through interacting with selective autophagy receptors. Unlike most ATG proteins, the ATG8 gene family is significantly expanded in vascular plants, but its functional specialization remains poorly understood. Using transcriptional and translational reporters in *Arabidopsis thaliana*, we revealed that ATG8 isoforms are differentially expressed across tissues and form distinct autophagosomes within the same cell. To explore ATG8 specialization, we generated the nonuple *Δatg8* mutant lacking all nine ATG8 isoforms. The mutant displayed hypersensitivity to carbon and nitrogen starvation, coupled with defects in bulk and selective autophagy as shown by biochemical and ultrastructural analyses. Complementation experiments demonstrated that ATG8A could rescue both carbon and nitrogen starvation phenotypes, whereas ATG8H could only complement carbon starvation. Proximity labeling proteomics further identified isoform-specific interactors under nitrogen starvation, underscoring their functional divergence. These findings provide genetic evidence for functional specialization of ATG8 isoforms in plants and lay the foundation for investigating their roles in diverse cell types and stress conditions.

## Introduction

Autophagy is an evolutionarily conserved cellular quality control mechanism essential for maintaining homeostasis and adapting to environmental stresses (*1–3*). It functions by selectively degrading and recycling damaged, redundant, or harmful cellular components, ensuring cellular integrity and energy balance (*4–6*). Although autophagy occurs constitutively, it is highly inducible under stress conditions such as nutrient deprivation, hypoxia, or infection (*7–10*). During these challenges, autophagy sustains survival by facilitating the degradation of intracellular material, which is sequestered into specialized double-membrane compartments called autophagosomes (*11, 12*). These structures subsequently fuse with lytic organelles—the vacuole in plants and yeast or the lysosome in animals—where their contents are broken down and recycled (*3*).

Contrary to initial views of autophagy as a non-selective bulk degradation process, it is now recognized as a highly selective pathway (*13–16*). This selectivity is mediated by specific interactions between cargo receptors, known as selective autophagy receptors (SARs), and autophagy-related proteins such as ATG8 (*17, 18*). These interactions enable the precise targeting of a wide range of substrates, from protein aggregates to damaged organelles, thus tailoring autophagic responses to specific cellular needs (*19*).

Autophagosome biogenesis progresses through three tightly regulated stages: initiation, expansion, and maturation. This process is orchestrated by the autophagy-related (ATG) protein family, comprising approximately 40 conserved members (*20, 21*). Central to autophagy is ATG8, a ubiquitin-like protein crucial for autophagosome formation, cargo recruitment, and membrane trafficking (*22, 23*). Once processed and lipidated, ATG8 associates with the autophagosome membrane, acting as a scaffold for the assembly of other core autophagy machinery and cargo receptors (*24, 25*).

Interestingly, unlike most ATG proteins, which exist as single or a few isoforms, the ATG8 gene family has undergone significant expansion in vascular plants (*26, 27*). While yeast and eukaryotes from early branching groups encode a single ATG8, vascular plants possess multiple isoforms, with *Arabidopsis thaliana* containing nine distinct ATG8 genes (AtATG8a-i) forming two major clades. Clade I contains ATG8A-G, whereas Clade II contains ATG8H and ATG8I (*28*). Despite moderate sequence divergence and differential expression patterns (*29, 30*)the biological significance of ATG8 isoform expansion and its implications for selective autophagy remain poorly understood.

Here, we present genetic evidence for the functional specialization of ATG8 isoforms in *A. thaliana* by generating a nonuple ATG8 mutant lacking all nine ATG8 genes. Our study reveals that ATG8 isoforms not only exhibit distinct tissue-specific expression and subcellular localization but also differ in their ability to mediate autophagic responses under specific stress conditions. These findings highlight the complex regulatory landscape of autophagy in plants and provide a foundation for unraveling the mechanisms underlying ATG8 specialization.

## Results and Discussion

### *Arabidopsis thaliana* ATG8 isoforms exhibit tissue-specific expression patterns and form distinct autophagosomes within root cells

To explore ATG8 specialization, we first checked the expression patterns of all the nine Arabidopsis ATG8 isoforms. We generated ATG8 promoter-GFP-GUS (pATG8X::GFP-GUS) expressing lines and performed β-glucuronidase (GUS) staining (Fig. 1A). Interestingly, AtATG8s exhibit distinct expression patterns and we only observed a partial overlap between different isoforms, indicating a certain degree of tissue specificity. ATG8E, ATG8F and ATG8G, show a more widespread expression pattern over different tissues and organs, whereas other isoforms, including ATG8A, ATG8B and ATG8I, appear to be restricted to the root (Fig. 1A). Furthermore, some isoforms exhibit peculiar tissue- or cell-type specificities, such as ATG8D being strongly induced in the apex of the cotyledon or ATG8C appearing specifically expressed in guard cells (Fig. 1A). These observations hint at a potential tissue- or cell-type-specific function of the different ATG8 isoforms.

**Figure 1.**
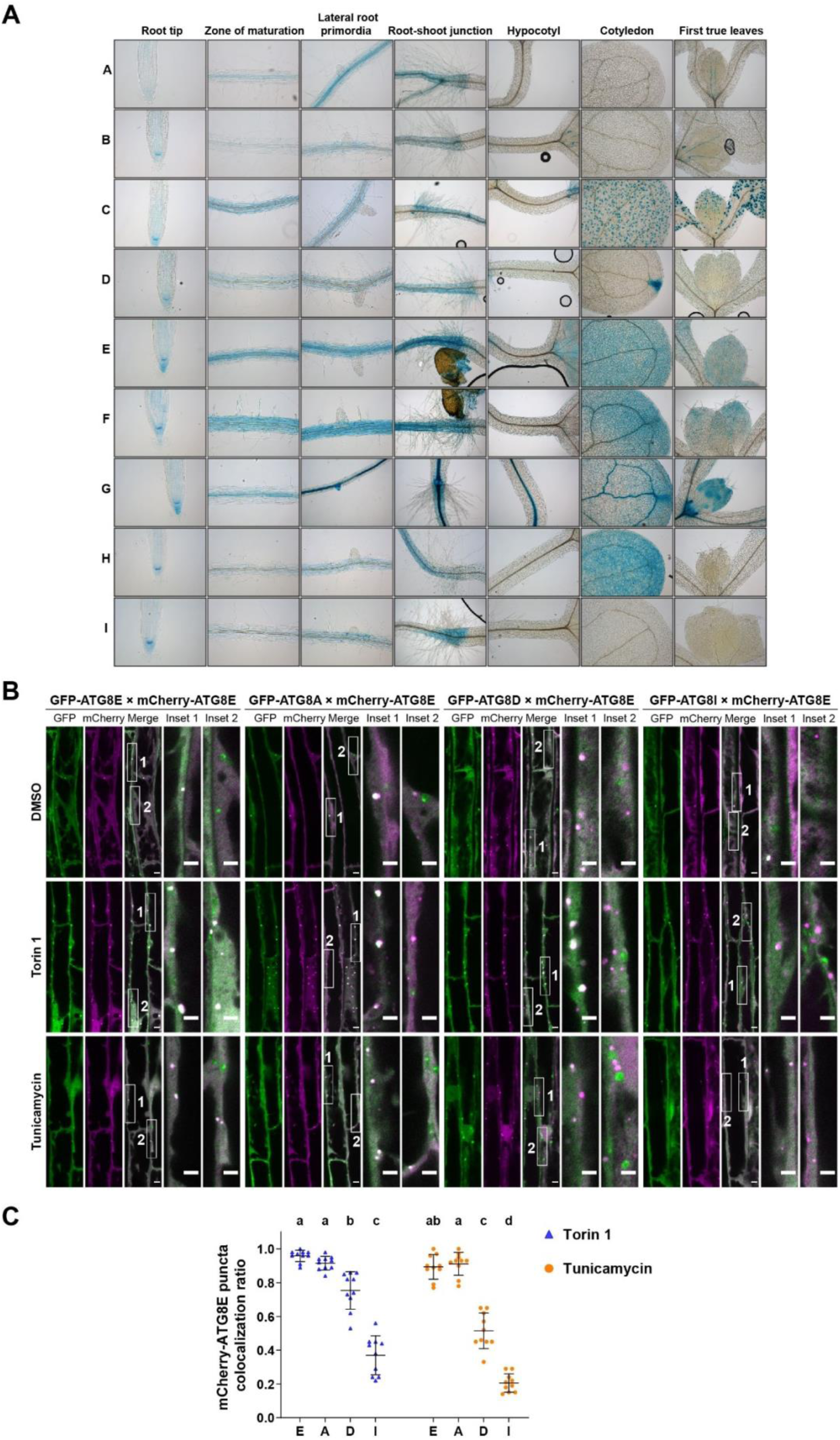
*Arabidopsis thaliana* ATG8 isoforms exhibit tissue-specific expression patterns and form distinct autophagosomes within root cells. (A) Representative GUS staining images showing the spatial-temporal expression patterns of 9 Arabidopsis ATG8 isoforms.10-days old Arabidopsis seedlings expressing pATG8X::GFP-GUS (X represents the 9 ATG8 isoforms, from A to I) were stained with GUS staining buffer. (B) Representative confocal microscopic images showing the colocalization of mCherry-ATG8E with different GFP-ATG8 isoforms in Arabidopsis root epidermal cells. 5-days old Arabidopsis seedlings co-expressing mCherry-ATG8E with GFP-ATG8E, GFP-ATG8A, GFP-ATG8D, or GFP-ATG8I were incubated in ½ MS liquid media containing either DMSO (as mock condition) for 2 h, 5 μM Torin1 for 2 h, or 10 μg/mL Tunicamycin for 4 h. Representative images of 10 replicates were shown here. Scale bars, 5μm. Inset scale bars, 3 μm. (C) Quantification of mCherry-ATG8E colocalization ratio of the Arabidopsis root epidermal cells imaged in (B). The mCherry-ATG8E colocalization ratio was calculated as the ratio of the number of mCherry-ATG8E puncta that colocalized with GFP-ATG8 isoforms to the total number of mCherry-ATG8E puncta. Bars indicate the mean ± SD of 10 replicates. Brown-Forsy and Welch ANOVA tests with Dunnett’s T3 multiple comparisons tests were used for statistically comparing the colocalization difference between each treatment group.

Then, we decided to test the subcellular compartmentalization of ATG8 isoforms. We co-expressed mCherry-ATG8E translational fusion constructs with GFP-tagged versions of ATG8A, ATG8D and ATG8I and assessed their co-localization upon bulk autophagy inducing chemical Torin1 (*31*) and ER-stress inducer Tunicamycin (*32*) treatments (Fig. 1B and C). Irrespective of the treatment, ATG8E colocalized almost entirely with ATG8A. Conversely, ATG8D and ATG8I exhibit a lower degree of colocalization with ATG8E during Torin 1 treatment and even more pronouncedly upon Tunicamycin treatment. ATG8I, representative of the clade II of ATG8 isoforms, has the weakest level of colocalization with ATG8E. These data indicate there are distinct pools of autophagosomes that are labelled with different ATG8 isoforms. In sum, the expression site polymorphism as well as varying levels of colocalization upon different treatments, support the hypothesis that ATG8 isoforms might fulfill different functions or respond to different stimuli.

### *atg8* nonuple mutant (*Δatg8*) is deficient in autophagy

The partial overlap in expression and colocalization patterns of ATG8 isoforms prompted us to generate an ATG8-free *Arabidopsis thaliana* line that we could use as a tool to investigate ATG8 specialization. Using multiplex CRISPR mutagenesis, we combined 9 guide RNAs (one for each ATG8 isoform) in a construct containing an intronized Cas9 (*33*) and transformed it to generate a nonuple knock-out of *atg8.* We first confirmed the mutations using whole genome sequencing (Fig. 2A). To functionally verify *Δatg8* as an autophagy deficient mutant, we performed the typical nutrient starvation assays (*34*). We transferred 9-days old Arabidopsis seedlings to carbon or nitrogen deprived ½ MS liquid medium for 4 or 6 days, respectively. Similar to the autophagy-deficient mutants *atg2* and *atg5* (*35, 36*), *Δatg8* exhibits reduced growth and discoloration of the cotyledons (Fig. 2B and C). To biochemically validate *Δatg8* mutant, we performed autophagic flux assays under carbon and nitrogen starvation conditions, by measuring the endogenous levels of the stereotypical autophagy substrate NBR1 (Fig. 2D andE) (*37*). NBR1 accumulated at a comparable level in both *Δatg8* and *atg5* under all conditions and was insensitive to concanamycin A treatment that blocks vacuolar degradation (*38*), denoting that both mutants are defective in autophagic degradation (Fig. 2D and E). Collectively, these results suggest that atg8 nonuple mutant is deficient in autophagy.

**Figure 2.**
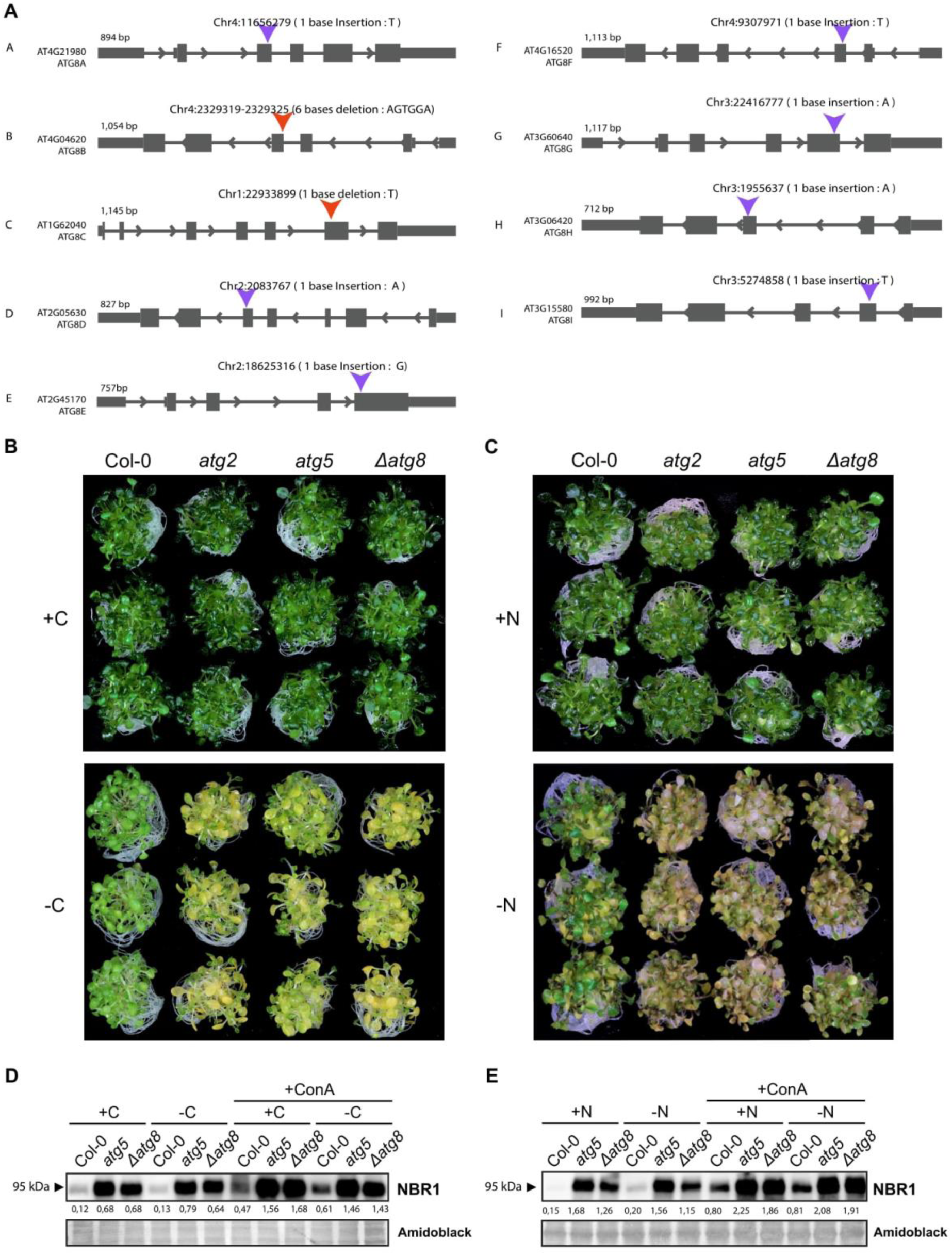
*atg8* nonuple mutant (*Δatg8*) is deficient in autophagy. (A) Schematic representation of gene models illustrating CRISPR-induced mutations in the ATG8 gene. Blue arrows denote insertion sites, while red arrows indicate deletion sites. The forward and reverse arrows within the gene model indicate the directionality of gene transcription, reflecting the strandedness of the gene. (B, C) Carbon (B) and nitrogen (C) starvation phenotypic assays comparing Col-0, *atg5* and *Δatg8* (n = 3) in carbon-rich (+C) or carbon-deficient (-C) ½ MS liquid medium and nitrogen-rich (+N) or nitrogen-deficient (-N) ½ MS liquid medium. (D, E) Western blots comparing endogenous NBR1 levels in Col-0, *atg5* and *Δatg8* upon carbon (D) and nitrogen (E) starvation, in combination with Concanamycin A (1μM). Relative quantification of protein bands is reported below the blots.

### *Δatg8* mutant is defective in selective autophagy

Nitrogen and carbon starvation are considered to trigger bulk autophagy (*39, 40*). To test if *Δatg8* is also defective in selective autophagy, we tested its ability to perform mitophagy and pexophagy. For mitophagy, we measured the levels of the outer mitochondrial membrane voltage dependent anion channel I (VDAC) and the matrix protein isocitrate dehydrogenase (IDH) upon 2,4-dinitrophenol (DNP) treatment. DNP is an uncoupler that leads to mitochondrial depolarization and triggers mitophagy (*41, 42*) (Fig. 3A). Although both VDAC and IDH levels were decreased in the DNP treated wild type Col-0 plants, *Δatg8* behaved similar to *atg5* and showed no change in VDAC or IDH levels (*41*). Likewise, when we assessed peroxisome degradation using the catalase antibody, *Δatg8* mutant behaved similar to *atg5* mutant and was unable to perform pexophagy (*43*) (Fig. 3B). Altogether, these results suggest *Δatg8* mutant is defective in mitophagy and pexophagy.

**Figure 3.**
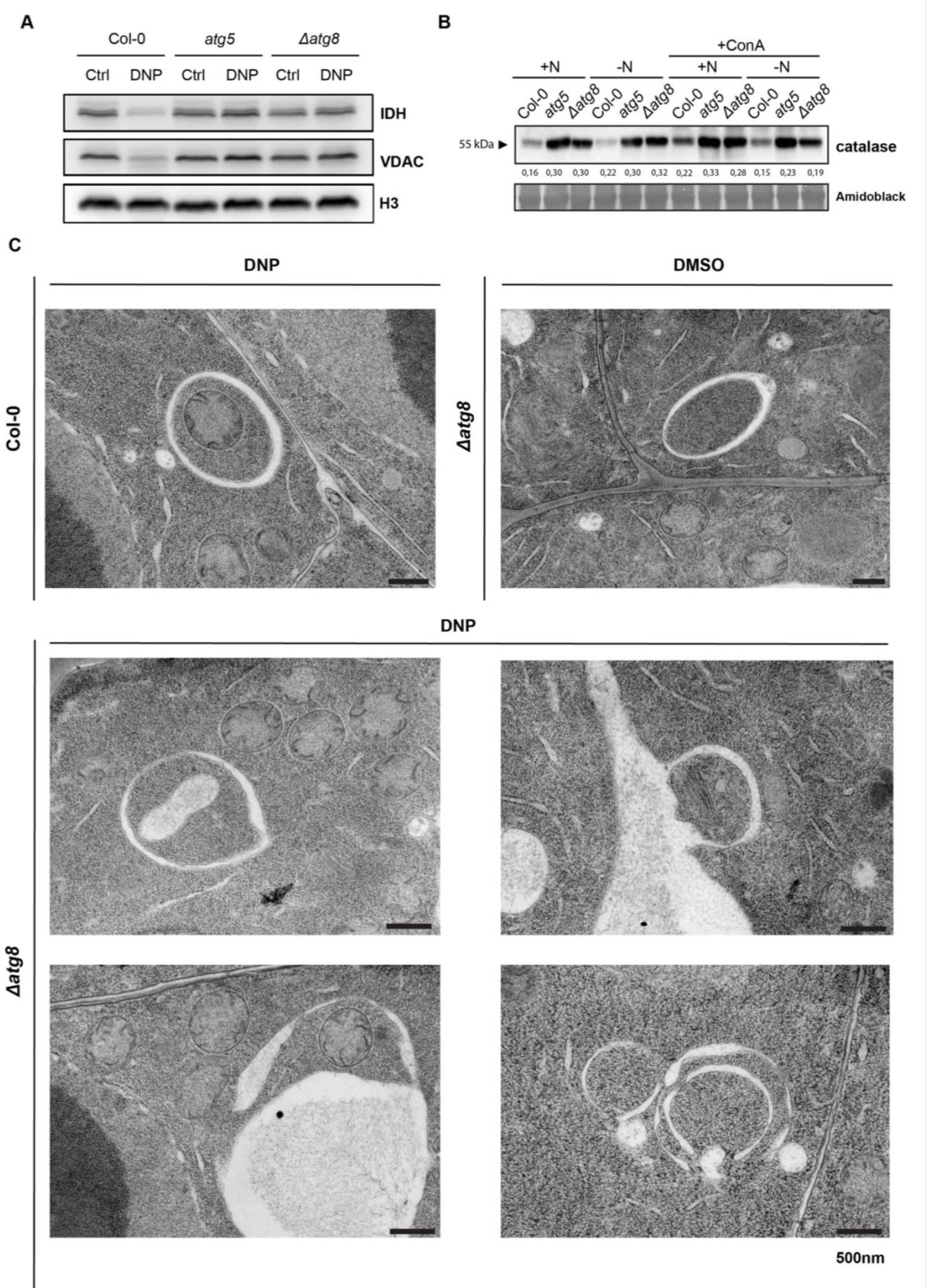
The *Δatg8* mutant is not able to perform mitophagy and pexophagy. (A) Western blots comparing endogenous IDH and VDAC levels in Col-0, *atg5* and *Δatg8* upon DNP treatment. (B) Western blots comparing endogenous catalase levels in Col-0, *atg5* and *Δatg8* upon N starvation treatment. Relative quantification of protein bands is reported below the blot.(C) Electron micrographs of Col-0 and *Δatg8* root cells treated with DNP or DMSO. Scale bars, 500 nm.

Next, we performed transmission electron microscopy (TEM) analysis of mitophagy in Arabidopsis root cells. Although we could detect mitophagosomes in wild type Col-0 plants, we did not observe any mitophagosomes in *Δatg8* mutant (Fig. 3C). However, we could still detect double-membraned structures that resemble autophagosomes in the *Δatg8* mutant. These vesicles appear to non-specifically engulf various types of cellular components. Although further studies are necessary to understand the nature of these compartments, a plausible source could be provacuoles, compartments that appear during vacuole biogenesis (*44*). Indeed, provacuole formation has been reported to be independent of autophagy as autophagy-deficient mutants can still form provacuoles (*44*). In summary, our data suggest *Δatg8* mutant is unable to carry out autophagic recycling.

### Complementation of *Δatg8* with ATG8A or ATG8H reveals functional specialization

After confirming the autophagy-deficient phenotype of the *Δatg8* mutant, to assess the functional specialization of ATG8 isoforms, we complemented it with GFP tagged ATG8A and ATG8H, representing both clades. First, we analyzed the complemented lines with confocal microscopy. Both GFP-ATG8A and GFP-ATG8H formed cytoplasmic bright puncta, which further accumulated in the vacuole upon Concanamycin A treatment and underwent autophagic flux (Fig. S2A-B).

To assess to what extent they were able to recover the autophagic function and evaluate potential isoform-specific responses to different stressors, we subjected the two complementation lines to starvation assays. Upon carbon starvation, both complementation mutants exhibit a similar phenotype to Col-0 (Fig. 4A). This suggests that both ATG8A and ATG8H could mediate the autophagic recycling of cellular material that ensure survival during carbon deprivation. In contrast, during nitrogen starvation only GFP-ATG8A expression recovered the sensitivity to nitrogen starvation. GFP-ATG8H expressing lines were similar to the *Δatg8* mutant (Fig. 4B). These results provide functional genetic evidence for ATG8H specialization in *Arabidopsis thaliana*.

**Figure 4.**
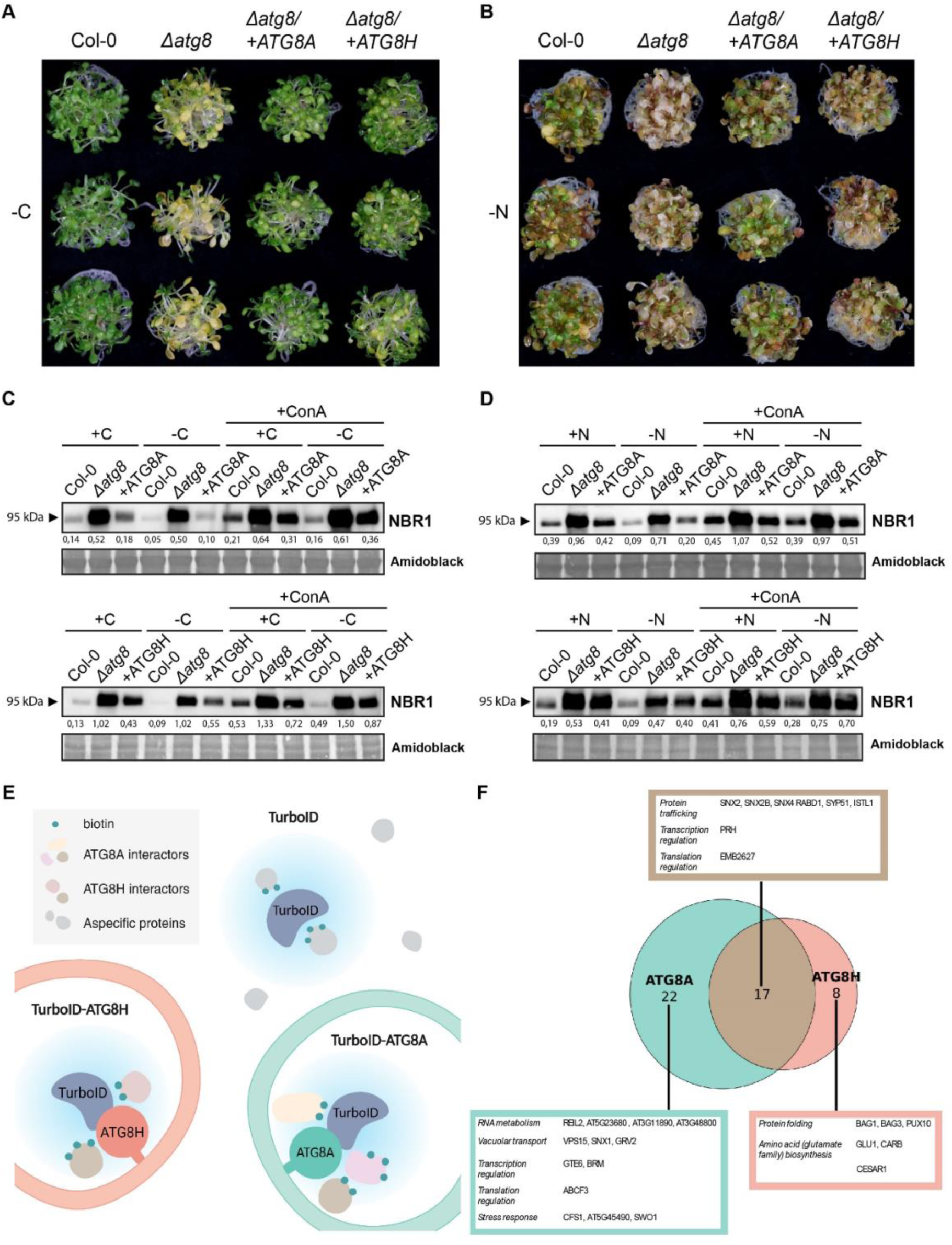
Complementation of *Δatg8* with ATG8A or ATG8H reveals functional specialization of ATG8 isoforms. (A, B) Carbon (A) and nitrogen (B) starvation phenotypic assays comparing Col-0, *Δatg8* and complementation lines *Δatg8* /+ATG8A, *Δatg8*/+ATG8H (n = 3) in carbon-deficient (-C) ½ MS liquid medium and nitrogen-deficient (-N) ½ MS liquid medium. (C, D) Western blots comparing endogenous NBR1 levels in Col-0, *Δatg8, Δatg8* /+ATG8A and *Δatg8* /+ATG8H upon carbon (C) and nitrogen (D) starvation, in combination with Concanamycin A (1μM). Relative quantification of protein bands is reported below the blots. (E) Schematic representation of TurboID proximity labeling analysis. (F) Venn diagram reporting common and unique interactors of ATG8A and ATG8H under nitrogen starvation.

To support these findings, we performed autophagic flux assays under carbon and nitrogen starvation. Under carbon starvation conditions, both *Δatg8/+GFP-ATG8A* and *Δatg8/+GFP-ATG8H* had similar NBR1 flux, in contrast to the *Δatg8* mutant (Fig. 4C). This is consistent with the phenotyping results and indicates that both ATG8A and ATG8H are able to trigger bulk autophagy in response to carbon deprivation. However, during nitrogen starvation NBR1 flux in *Δatg8/+GFP-ATG8H* was similar to the *Δatg8* mutant (Fig. 4D), further corroborating the hypothesis that ATG8H, unlike ATG8A, is not able to fully operate autophagy in response to nitrogen deprivation.

### ATG8A and ATG8H have distinct proxitomes during nitrogen starvation

Following the observation that ATG8A and ATG8H do not respond equally to N starvation stress, we reasoned that the two isoforms may be interacting with different proteins that are involved in autophagy signaling or cargo recognition. Indeed, our results could be explained by the inability of ATG8H to engage with the nitrogen starvation response signaling, or its failure to associate with the selective autophagy receptors recognizing the cargoes that need to be degraded to cope with nitrogen deprivation. To test these hypotheses, we complemented the *Δatg8* mutant with the biotinylating enzyme TurboID fused ATG8A and ATG8H and determined the ATG8 proxitomes during nitrogen starvation (Fig. 4E). We used TurboID alone as negative control. 47 proteins exhibited specific association with both or only one of the two ATG8 isoforms (Supplementary Table 3). Among these, numerous well-known interactors were found, including several ATG proteins. Interestingly, whereas some ATG proteins showed similar levels of association with ATG8A and ATG8H, such as ATG3, ATG7, ATG14B and ATG18F, other ATG proteins appear to interact prevalently or exclusively with just one isoform (Supplementary Table 3). In the case of ATG1, ATG1A seems to interact with ATG8A uniquely, while ATG1B can associate with both isoforms but still exhibits a stronger association with ATG8A in the conditions tested. ATG1 kinase initiates autophagosome biogenesis and constitutes one of the major targets for autophagy regulation (*45, 46*). In light of this, our results may indicate that during N starvation ATG1A and ATG1B recruit ATG8A preferentially to promote nutrient replenishment. It is also interesting to observe that less than one third of the total ATG8 interactors are shared between ATG8A and ATG8H (Fig. 4F), whilst 22 proteins specifically interact with ATG8A, and 8 proteins interact with only ATG8H. As a proof on concept, our proximity labeling analysis suggested that the adaptor protein CFS1 specifically interacts with ATG8A but not with ATG8H, consistent with our previous results (*47*). While other interactors need to be further validated, these results suggest single-isoform TurboID lines provide an effective tool to study ATG8 specialization in a wide range of stress conditions.

Since the core autophagy machinery is shared across various selective autophagy pathways, the question of how cells achieve subcellular compartmentalization of these concurrent mechanisms remains unresolved. A plausible explanation is ATG8 isoform specialization, whereby distinct ATG8 variants interact with specific adaptors, receptors, or ATG proteins to direct and compartmentalize autophagic processes. Previous biochemical and proteomic analyses in potato supported this hypothesis, revealing isoform-specific interactomes (*48*). However, the co-occurrence of multiple ATG8 isoforms within individual autophagosomes (Fig. 1) suggests that some unique interactors might have been overlooked.

In this study, we provide genetic evidence for ATG8 specialization in plants using an Arabidopsis *Δatg8* nonuple mutant complemented with individual ATG8 isoforms. Unlike mitophagy observed in HeLa cells lacking multiple ATG8 genes, the Arabidopsis *Δatg8* mutant failed to perform mitophagy and pexophagy, as evidenced by the absence of mitophagosomes and the accumulation of autophagy substrates (Fig. 3). While double-membraned vesicles were observed in *Δatg8* cells, these are likely provacuolar compartments, which are independent of autophagy.

Interestingly, complementation experiments revealed functional divergence among ATG8 isoforms. While both ATG8A and ATG8H restored carbon starvation sensitivity, only ATG8A was able to complement nitrogen starvation sensitivity (Fig. 4). Consistent with their specialization, proximity labeling proteomics demonstrated distinct interactomes for ATG8A and ATG8H. Further studies are necessary to link the nitrogen sensitivity phenotype to the differentially interacting proteins. Nevertheless, our findings establish the functional specialization of ATG8 isoforms in plants, providing a framework for understanding how cells fine-tune autophagic processes in response to diverse and overlapping signals.

## Material and Methods

### Plant material and cloning

All Arabidopsis thaliana lines used originate from the Columbia (Col-0) ecotype. The *Δatg8* mutant was generated with two rounds of CRISPR editing. First, the CRISPR/Cas9 construct including gRNAs for all nine AtATG8 genes and an intronized Cas9 was assembled according to a protocol previously described (*33*). The construct was assembled in binary vector pAGM62636 (*33*) that contains a p15 origin of replication for low copy number replication in *E. coli* and an *Agrobacterium rhizogenes* A4 ori for single copy replication in Agrobacterium, resulting in plasmid pAGM70811. This vector backbone was chosen to minimize the risk or recombination between the 9 guides RNA cassettes present on the same plasmid. The gRNAs sequences are the following:

ATG8A (AT4G21980): AGCTTACGGGAATTCTGTCA
ATG8B (AT4G04620): GAACTCAATACAGGTGATTG
ATG8C (AT1G62040): CAGTTAGATCAGCTGGAACA
ATG8D (AT2G05630): AAGAGGATGTTCATGCTTG
ATG8E (AT2G45170): CTGTTAGGTCTGATGGCACA
ATG8F (AT4G16520): TTCAGAGAAGAGAAGGGCAG
ATG8G (AT3G60640): GAGGAGACAGTACCGGTGGG
ATG8H (AT3G06420): AAACGCAGATCTGCCAGACA
ATG8I (AT3G15580): GATGAAAGGCTCGCGGAGTCG

Primary transformants were genotyped by amplicon sequencing. The data was analysed using a custom-built pipeline to retrieve indel frequencies in the ATG8 genes, using bwa and samtools *mpileup* (*49, 50*). Code and documentation are available at https://github.com/pierre-bourguet/CRISPR_genotyping. Following sequencing, we could only confirm the successful mutation of eight of the nine isoforms. Therefore, a second CRISPR/Cas9 was employed to target the last gene (ATG8D – AT2G05630). The construct was assembled according to the protocol previously described (*51*), on pHEE401E plasmid, with the following gRNAs sequences: GAACAACAGAGACTCGACCA, GTGATGTCCCGGATATTGAT. The nonuple mutant *Δatg8* contains both T-DNA CRISPR/Cas9 cassettes and is resistant to BASTA and Hygromycin. The mutations of nine ATG8 isoforms were confirmed via single cell sequencing.

All the plasmids, except pAGM7081, were assembled through the GreenGate cloning procedure (*52*) and were constructed as follows: pGGZ003_ATG8X::GFP-GUS (X represents the 9 ATG8 isoforms, from A to I), pGGSUN_RPS5::mCherry-TurboID; pGGSUN_RPS5::mCherry-TurboID-ATG8A; pGGSUN_RPS5::mCherry-TurboID-ATG8H; pGGSUN_HTR5::GFP-ATG8A; pGGSUN_HTR5::GFP-ATG8H. Apart from pATG8X::GFP-GUS expressing plants, which are hygromycin resistant, transformants were selected via seed coat fluorescence. The coding sequences of ATG8A and ATG8H carry silent mutations to avoid CRISPR/Cas9 targeting. The point mutations from the start codon are the following: ATG8A, 81bp T > A, 84bp C > T, 87bp A > G, 93bp C > G; for ATG8H, 123bp C > T, 126bp A > C, 129bp T > C, 135bp A > G, 138bp C > T.

### DNA sequencing and analysis

High-quality DNA was extracted using the cetyltrimethylammonium bromide (CTAB) method and used to construct Illumina-compatible libraries with the Nextera XT DNA Library Preparation Kit, following the manufacturer’s instructions. Sequencing was performed on an Illumina NextSeq instrument in paired-end 150 bp mode.

Raw FASTQ files obtained from sequencing were quality-checked and adapter-trimmed using TrimGalore (https://github.com/FelixKrueger/TrimGalore) with default settings. The trimmed FASTQ files were aligned to the TAIR10 genome using Bowtie2 (*51*) with the parameters `-D 15 - R 2 -N 0 -L 22 -i S,1,1.15`. The resulting aligned BAM files were sorted and indexed using SAMtools (*52*) and manually inspected for insertions or deletions using the Integrative Genomics Viewer (IGV). Deletions and insertions were manually inspected, and the corresponding changes in cDNA and protein sequences were catalogued (Supplementary Table 1).

### Plant growth and treatments

For standard plant growth, Arabidopsis seeds were gas sterilized with sodium hypochlorite + HCl (10:1 v/v), sown on water-saturated soil and grown in 16h light/8h dark photoperiod with 165 μmol m^-2 s^-1 light intensity. For *in vitro* growth, Arabidopsis seeds were surface sterilized in 70% ethanol for 10 minutes twice, then rinsed in absolute ethanol and dried on sterile paper. Seeds were sown in ½ MS liquid medium (Murashige and Skoog salt + Gamborg B5 vitamin mixture [Duchefa] supplemented with 0.5 g/liter MES and 1% sucrose, pH 5.7), vernalized at 4 °C in the dark for 2 days, and then grown under LEDs with 85μM/m²/s with a 14 h light/10 h dark photoperiod.

For drug treatments, all drugs were dissolved in DMSO and added to the desired concentration: 3 µM Torin 1 (Santa Cruz Biotechnology – CAS 1222998-36-8), 10 µg/mL Tunicamycin (Santa Cruz Biotechnology – CAS 11089-65-9), 1-2 µM Concanamycin A (Santa Cruz Biotechnology – CAS 80890-47-7), 50 µM DNP (Sigma-Aldrich, D198501-1KG). An equal amount of pure DMSO was added to control samples.

For confocal microscopy, Arabidopsis seeds were sterilized by 70% ethanol + 0.05% Tween 20 (Sigma-Aldrich) for 5 min and were subsequently sterilized by 100% ethanol for 10 min. Sterilized seeds were stored in sterile water at 4°C for 1 d for vernalization. Vernalized seeds were spread on 1/2 MS media plates (+1% plant agar [Duchefa]) and vertically grown at 21°C at 60% humidity under LEDs with 50 mM/m^2^s a and a 16 h light/8 h dark photoperiod for 5 days. 5-days old seedlings were incubated in ½ MS media containing either DMSO for 2 h, 3 μM Torin 1 for 2 h, 10 μg/mL Tunicamycin for 4 h or 2 μM Concanamycin for 2.5 h before imaging.

### Carbon and Nitrogen starvation assays

*A. thaliana* seeds (∼30 per sample, 3 replicates per condition) were sterilized with ethanol, sown in ½ MS liquid medium, vernalized at 4 °C in the dark for 2 days and grown at 21 °C under LEDs with 85μM/m2/s and a 14 h light/10 h dark photoperiod. The starvation treatments were performed on 9-days old seedlings, by replacing ½ MS liquid medium with the same medium (as control), ½ MS liquid medium without sucrose for C starvation or ½ MS liquid medium without nitrogen (Murashige & Skoog without Nitrogen, Caisson Laboratories – MSP21) for N starvation. Prior to medium replacement, the seedlings were washed twice with 1 mL of the new medium to ensure proper removal of the previous medium. For C starvation, the seedlings were kept in the dark. Pictures of the samples were taken after 4 days of C starvation and 6 days of N starvation.

### Confocal microscopy

For confocal microscopy, Arabidopsis seedlings were placed on a microscope slide with water and covered with a coverslip. The epidermal cells of root transition and elongation zone were used for image acquisition.

Confocal images were acquired via an upright point laser scanning confocal microscope ZEISS LSM800 Axio Imager.Z2 (Carl Zeiss) equipped with high-sensitive GaAsP detectors (Gallium Arsenide), a LD C-Apochromat 40X objective lens (numerical aperture 1.1, water immersion), and ZEN software (blue edition 3.8, Carl Zeiss). GFP signals were excited at 488 nm and detected between 488 and 545 nm. mCherry signals were excited at 561 nm and detected between 570 and 617 nm.

### Image processing and statistics

Confocal images were processed and quantified by Fiji (version 1.52, Fiji). The mCherry-ATG8E colocalization ratio was calculated as the ratio of the number of mCherry-ATG8E puncta that colocalized with GFP-ATG8 isoforms to the total number of mCherry-ATG8E puncta. Statistics tests were performed via GraphPad Prism 8.1.1.

### Protein extraction and western blotting

20-40 *A. thaliana* seeds per sample were surface sterilized with ethanol, sown in ½ MS liquid medium, vernalized at 4 °C in the dark for 2 days and grown for 7 days at 21 °C under LEDs with 85μM/m2/s with a 14 h light/10 h dark photoperiod. For starvation treatments, performed overnight, the ½ MS liquid medium was replaced with the same medium (as control), ½ MS liquid medium without sucrose for C starvation or ½ MS liquid medium without nitrogen (Murashige & Skoog without Nitrogen, Caisson Laboratories – MSP21) for N starvation. For C starvation, the samples were kept in the dark. When required, 1 µM Concanamycin A (Santa Cruz Biotechnology – CAS 80890-47-7) was added to the new medium. The seedlings were harvested in safe lock Eppendorf tubes containing 2 mm Ø glass bead, flash frozen in liquid nitrogen and pulverized using a Silamat S7 (Ivoclar vivident). Total proteins were extracted in 2X Laemmli buffer by agitating the samples in the Silamat S7 for 20 s. The samples were boiled at 70 °C and 1000 rpm shaking for 10 min, then centrifuged at max speed with a benchtop centrifuge. Total proteins were quantified with the amido black method. 10 µl of supernatant was added to 190 µl of deionized water, vortexed and then mixed with 1 ml of Amido Black Buffer (10% acetic acid, 90% methanol, 0.05% [w/v] Amido Black (Napthol Blue Black, Sigma N3393)) by inverting the tubes. After 10 min centrifugation at max speed, pellets were washed with 1 ml of Wash Buffer (10% acetic acid, 90% ethanol), mixed by inversion, and centrifuged for another 10 minutes at max speed. Pellets were resuspended in 0.2N NaOH and OD_630 nm_ was measured, with NaOH solution as blank, to quantify protein concentration with the OD = a[C] + b determined curve. 15 µg of total protein extracts were separated on SDS-PAGE gels and blotted onto PVDF Immobilon-P membrane (Millipore). NBR1 was detected using the anti-NBR1 antibody (Rabbit polyclonal; Agrisera – AS14 2805) diluted 1:10000. Catalase proteins were detected with Anti-Cat antibody (Rabbit polyclonal; Agrisera – AS09 501), diluted 1:1000. GFP was detected with the anti-GFP antibody (Mouse monoclonal; Roche – 11814460001), diluted 1:5000. Rabbit polyclonal antibody was detected with a goat anti-rabbit IgG HRP-linked antibody (Invitrogen, 65-6120) diluted 1:5000. Hybridized membranes were reacted with SuperSignal™ West Pico PLUS Chemiluminescent Substrate (Thermo Fisher Scientific) and imaged using an iBright CL1500 Imaging System (Invitrogen).

Protein bands were quantified with ImageJ according to the protocol previously described (*53*) and normalized on the loading control. The values reported on the figures correspond to mean values for 3 biological replicates.

For mitophagy assays, 5-days old Arabidopsis seedlings were treated with 50 µM DNP (Sigma-Aldrich, D198501-1KG), or an equal amount of DMSO, for 2-3 hours in the dark, then moved to liquid ½ MS medium for 1h recovery under light. Protein extraction and immunoblot analysis was performed as previously reported (*41*).

### GUS staining

10-days old Arabidopsis seedlings expressing pATG8X::GFP-GUS (X represents the 9 ATG8 isoforms, from A to I) were first immersed in acetone for 20 minutes and then washed with the GUS buffer (50 mM NaPO_4_, 2 mM K-ferrocyanide, 2mM K-ferricyanide, 0.2% Triton X-100). The washed samples were subsequently incubated in GUS staining buffer [GUS buffer + 2 mM X-Gluc (Thermo Scientific)] under 37℃ until a blue coloration was visible. The stained samples were then washed and discolored with 100% ethanol and were ready for photographing.

### Affinity purification of biotinylated proteins and nanoLC-MS/MS Analysis

*A. thaliana* seeds were surface sterilized with ethanol, stratified for 2 days at 4 °C in the dark and then grown in ½ MS (Duchefa)/0.5% MES/1% sucrose under LEDs with 85 μM/m2/s and a 14 h light/10 h dark photoperiod. 7-days old seedlings were washed and treated with N deficient ½ MS medium (or control ½ MS medium) overnight and the following morning 50 μM biotin was added to the medium. After 1 hour of biotin incubation, the seedlings were quickly rinsed in ice cold water, dried and frozen in liquid nitrogen. Around 1 gram of plant tissue was used for each sample and the affinity purification of biotinylated proteins was performed as previously described (*54*).

For MS Analysis, the nano HPLC system (UltiMate 3000 RSLC nano system) was coupled to an Orbitrap Exploris 480 mass spectrometer, equipped with a Nanospray Flex ion source (all parts Thermo Fisher Scientific). Peptides were loaded onto a trap column (PepMap Acclaim C18, 5 mm × 300 μm ID, 5 μm particles, 100 Å pore size, Thermo Fisher Scientific) at a flow rate of 25 μl/min using 0.1% TFA as mobile phase. After loading, the trap column was switched in line with the analytical column (PepMap Acclaim C18, 500 mm × 75 μm ID, 2 μm, 100 Å,Thermo Fisher Scientific). Peptides were eluted using a flow rate of 230 nl/min, starting with the mobile phases 98% A (0.1% formic acid in water) and 2% B (80% acetonitrile, 0.1% formic acid) and linearly increasing to 35% B over the next 120min. This was followed by a steep gradient to 95% B in 1 min, stayed there for 6 min and ramped down in 2 min to the starting conditions of 98% A and 2% B for equilibration at 30°C. The Orbitrap Exploris 480 mass spectrometer was operated in data-dependent mode, performing a full scan (m/z range 350-1200, resolution 60,000, normalized AGC target 300%) at 3 different compensation voltages (CV -45V, -60V and -75V), followed by MS/MS scans of the most abundant ions for a cycle time of 0.9 seconds for each. MS/MS spectra were acquired using an isolation width of 1.2 m/z, normalized AGC target 200%, HCD collision energy of 30 %, maximum injection time mode set to custom and resolution of 30,000. Precursor ions selected for fragmentation (include charge state 2-6) were excluded for 45 s. The monoisotopic precursor selection (MIPS) mode was set to peptide and the exclude isotopes feature was enabled.

### MS Data processing

For peptide identification, the RAW-files were loaded into Proteome Discoverer (version 2.5.0.400, Thermo Scientific). All MS/MS spectra were searched using MSAmanda v2.0.0.19924 (Dorfer V. et al., J. Proteome Res. 2014 Aug 1;13(8):3679-84). The peptide mass tolerance was set to ±10 ppm and fragment mass tolerance to±10 ppm, the maximum number of missed cleavages was set to 2, using tryptic enzymatic specificity without proline restriction. The RAW-files were searched against the Arabidopsis database (32,785 sequences; 14,482,855 residues), supplemented with common contaminants and sequences of tagged proteins of interest. The following search parameters were used: Oxidation on methionine, phosphorylation on serine, threonine, and tyrosine, deamidation on asparagine and glutamine, iodoacetamide derivative on cysteine, beta-methylthiolation on cysteine, biotinylation on lysine, ubiquitinylation residue on lysine, ubiquitination on lysine, pyro-glu from q on peptide N-terminal glutamine, acetylation on protein N-Terminus were set as variable modifications. The result was filtered to 1 % FDR on protein level using the Percolator algorithm (Käll L. et al., Nat. Methods. 2007 Nov; 4(11):923-5) integrated in Proteome Discoverer. The localization of the post-translational modification sites within the peptides was performed with the tool ptmRS, based on the tool phosphoRS (Taus T. et al., J. Proteome Res. 2011, 10, 5354-62). Additionally, an Amanda score cut-off of at least 150 was applied. Protein areas have been computed in IMP-apQuant (Doblmann J. et. al, J Proteome Res 2019, 18(1):535-41) by summing up unique and razor peptides. Resulting protein areas were normalized using iBAQ (Schwanhäusser B. et al., Nature 2011, 473(7347):337−42) and sum normalization was applied for normalization between samples. Match-between-runs (MBR) was applied for peptides with high confident peak area that were identified by MS/MS spectra in at least one run. Proteins were filtered to be identified by a minimum of 2 PSMs in at least 1 sample and quantified proteins were filtered to contain at least 3 quantified peptide groups. Statistical significance of differentially expressed proteins was determined using limma (Smyth, G. K. - Linear models and empirical Bayes methods for assessing differential expression in microarray experiments. Statistical Applications in Genetics and Molecular Biology, 2004, Volume 3, Article 3.) (Supplementary Table 2).

To identify potential interactors, log2FC was calculated comparing average PSMs in TID-ATG8A samples (n = 3) or TID-ATG8H samples (n = 3) against the TID control (n = 3). Proteins with average PSMs > 3, log2FC > 1 and p-value > 0,05 were selected as potential interactors. The list of potential interactors is provided in Supplementary Table 3.

### TEM

The TEM assay was performed following the previously established method (*41, 55*). Briefly, 5-days old *Arabidopsis* seedlings were germinated on ½ MS plates and dissected under microscopy before freezing. For high-pressure freezing, the root tips were collected and immediately frozen with a high-pressure freezer (EM ICE, Leica). For freeze substitution, the root tips were substituted with 2% osmium tetroxide in anhydrous acetone and maintained at −80 °C for 24 hours using an AFS2 temperature-controlling system (Leica). Subsequently, the samples were subjected to three washes with precooled acetone and slowly warmed up to room temperature over a 60-h period before being embedded in EPON resin. After resin polymerization, samples were mounted and trimmed. For the ultrastructure studies, 100 nm thin sections were prepared using an ultramicrotome (EM UC7, Lecia) and examined with a Hitachi H-7650 TEM (Hitachi-High Technologies) operated at 80 kV.

## Data availability

All the source data used to generate the main and supplementary figures are deposited to Zenodo (10.5281/zenodo.14277422). Genomic sequencing data generated in this study are deposited at the National Center for Biotechnology Information Gene Expression Omnibus (NCBI GEO, https://www.ncbi.nlm.nih.gov/geo/) under accession number GSE283481. The mass spectrometry proteomics data have been deposited to the ProteomeXchange Consortium via the PRIDE partner repository.

## Acknowledgements

We thank Vienna Biocenter Core Facilities (VBCF), particularly Proteomics, BioOptics and Plant Sciences. We thank the CLIP cluster (https://clip.science) for access for image analysis. We acknowledge funding from Austrian Academy of Sciences, Austrian Science Fund (FWF, P32355, P34944, SFB F79, DOC 111), Vienna Science and Technology Fund (WWTF, LS17-047, LS21-009), and European Research Council Grant (Project number: 101043370). Peng Gao is supported by the Vienna International Postdoctoral Program (VIP2) and Marie Curie Fellowship (Project number: 847548).

The authors declare no competing financial interest.

---

**Supplementary Figure 1.**
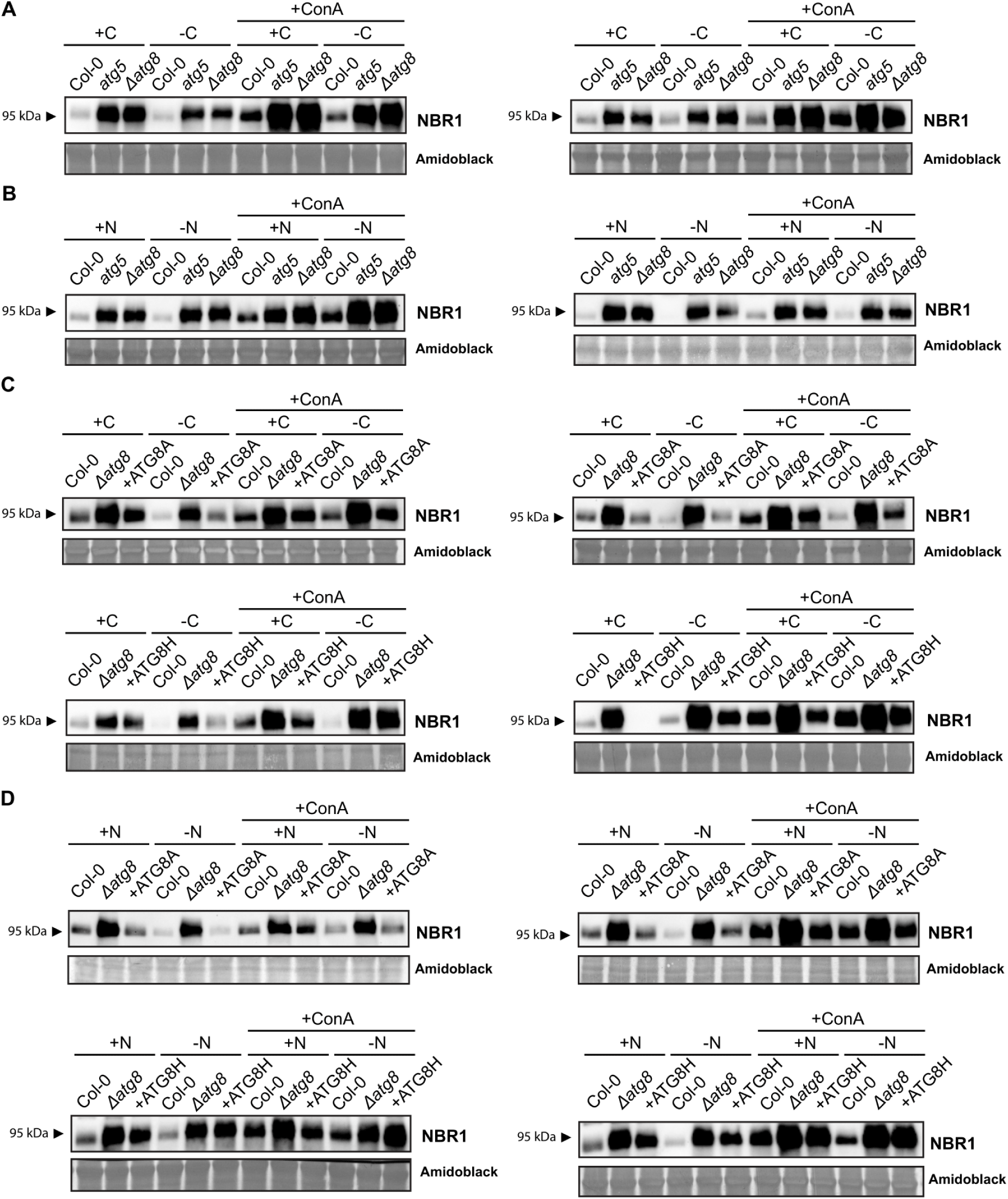
Replicates of western blots in Figure 2 (A, B) and Figure 4 (C, D).

**Supplementary Figure 2.**
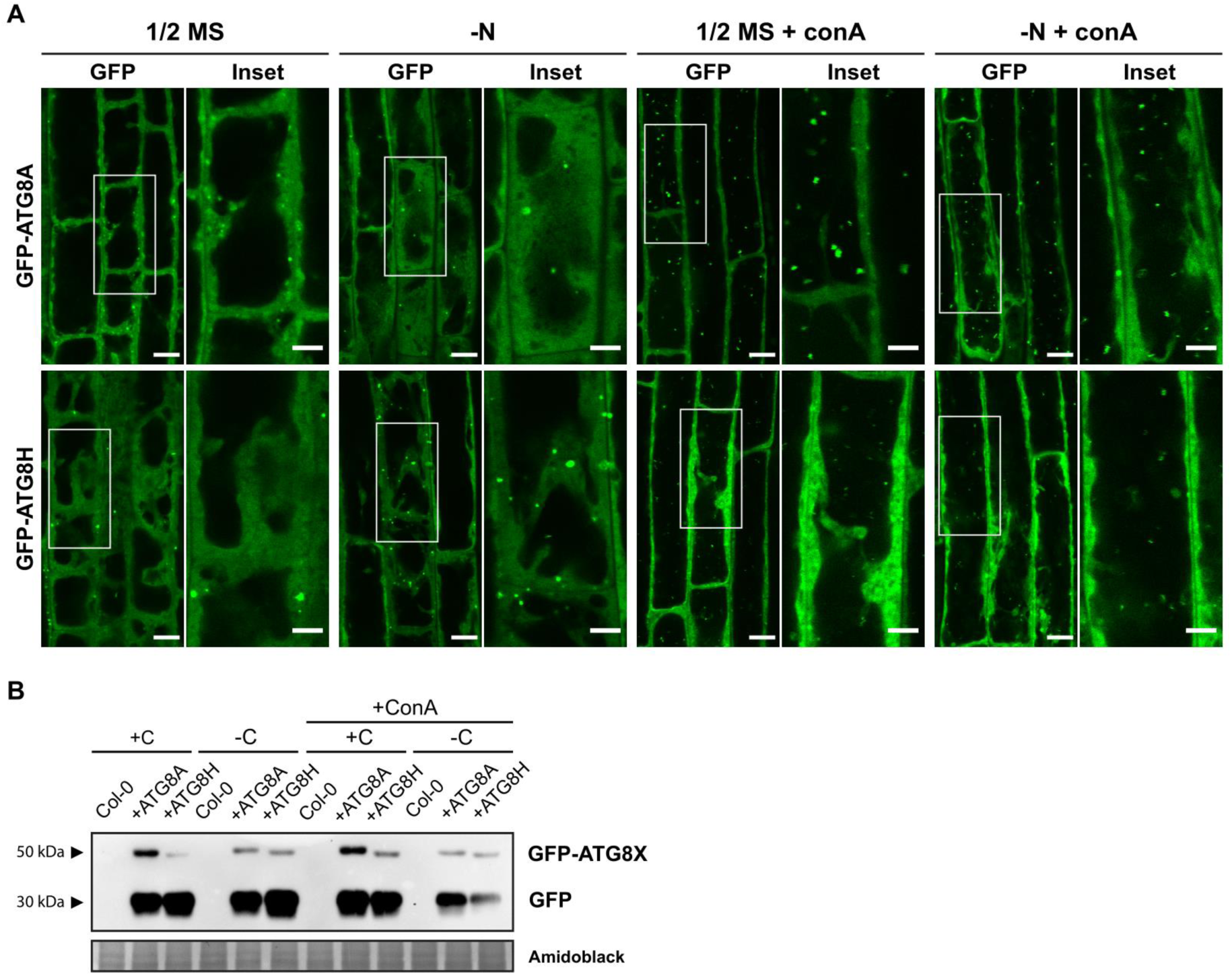
Arabidopsis ATG8 complementation lines have normal autophagosome structure and autophagic flux under control and nutrient-deficient conditions. (A) Representative confocal microscopic images showing the autophagosomes and the autophagic bodies inside the vacuole in root epidermal cells of the complementation lines Δatg8 /+GFP-ATG8A and Δatg8 /+GFP-ATG8A. 5-days old Arabidopsis seedlings were incubated in ½ MS liquid media or nitrogen-deficient (-N) liquid media for 3 h, or ½ MS liquid media or nitrogen-deficient (-N) liquid media containing 2 μM concanamycin A for 2.5 h before imaging. Representative images of 3 replicates were shown here. Scale bars, 10 μm. Inset scale bars, 5 μm. (B) Western blot comparing autophagic flux of complementation lines Δatg8 /+GFP-ATG8A and Δatg8 /+GFP-ATG8A upon C starvation treatment, in combination with concanamycin A (1μM).

**Supplementary Figure 3.**
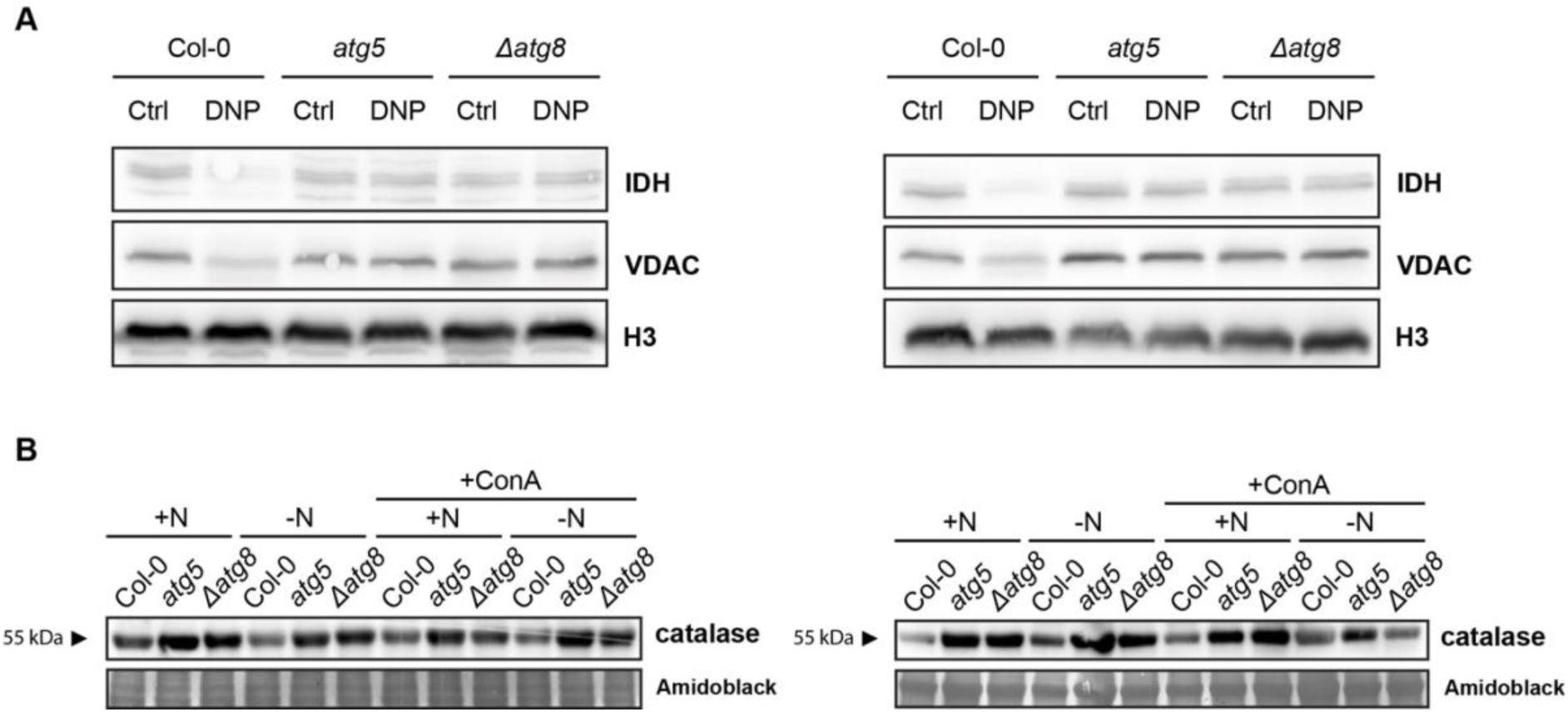
Replicates of western blots in Figure 3.

